# Mapping metabolic phases through online pressure and backscatter rates

**DOI:** 10.64898/2026.06.08.729528

**Authors:** Martin Malthe Borch, Pauline Kehr, Philip J. Gorter de Vries, Alex Toftgaard Nielsen

**Author notes:** Correspondence: Alex Toftgaard Nielsen.

## Abstract

Microbial metabolism can be represented as an energy-conserving process (catabolism) and a biomass-forming reaction (anabolism). Anabolism is traditionally measured through the turbidity of the culture, while catabolism is often assessed by the substrates consumed or the products formed. Standard measurements of biomass and products are intrusive and disrupt cultivation and headspace composition, potentially masking important analytical parameters and interactions.

Online pressure and backscatter were combined in small-scale closed batch vials to obtain undisturbed real-time measurements of catabolic and anabolic rates, enabling mapping of metabolic phases throughout an entire batch cultivation cycle.

The method identified discrete metabolic phases in yeast cultivation and thermophilic syngas fermentation. In nutrient-rich yeast cultivation, five metabolic phases were characterized, covering growth-associated and non-growth-associated gas formation. In a mixed community syngas fermentation, estimates of catabolic and anabolic rates distinguished early biomass increase from minimal net pressure change from two later gas-driven phases. An initial phase with a higher growth rate, linked to carboxydotrophy, followed by a phase with slightly lower growth and increased gas consumption, corresponding to hydrogenotrophic acetogenesis.

The study demonstrates that a simple, affordable experimental setup with online pressure and backscatter measurements can be used to visualize phase-plane mapping of microbial metabolism. An additional advantage is the ability to detect sequential metabolic cascades in mixed microbial communities, which is not possible with gas-sparging bioreactor studies. Using a single simple batch culture, growth and maintenance data can be obtained, even when growth is low or absent, thereby yielding parameters applicable to phenotypic characterization and dynamic metabolic modelling.

## 1 Introduction

A key method in the history of microbiology is the detection of bacterial activity through the detection of growth using turbidity (Koch 1961; Pasteur 1857). The continuous culture (chemostat) enabled a further characterization through identification of steady states at different substrate concentrations and dilution rates (D), leading to a model for microbial metabolism divided into a growth-associated and a non-growth-associated term (Novick and Szilard 1950; Monod 1949; Pirt 1965; Herbert 1959). Traditionally, chemostat measurement of product formation and substrate consumption is used as a measure of the catabolic activity, and to derive the maintenance model parameters (Villadsen et al. 2011; Heijnen and Kleerebezem 2010). When catabolism (product rates) and anabolism (growth rate) are measured simultaneously, microbial activity can be mapped onto a phase plane, visualising the parameters of the metabolic maintenance model. This concept was used by (Benthin et al. 1994; Villadsen et al. 2011) to plot product and growth rates on a phase plane. They characterised the metabolism of *Lactococcus cremoris* in batch and chemostat fermentations, revealing metabolic correlations and shifts with the changing growth conditions. But the maintenance model does not apply at low growth rates or at the single-cell level, because growth is enforced, and assumptions of steady states at the reactor level cannot be extrapolated to the single-cell level (Pirt 1987). Reviews highlight that the maintenance parameters should not be viewed as constants and are variables that change along the cultivation (Van Bodegom 2007). A chemostat-derived growth-enforced dataset will naturally favour growth-focused interpretations. Linear genome-scale modelling workflows based on chemostat data use enforced growth and constant maintenance parameters and extend them to the single-cell level. Naturally, if the models are fitted to growth-enforced chemostat data, the objective function that best fits the empirical data will be the one that optimises biomass growth. Though other objectives have been suggested, the most widespread objective function in metabolic modelling is growth optimisation (Feist and Palsson 2010). This idea is also presented by (Yasemi and Jolicoeur 2021) and they propose dynamic metabolic flux balance analysis models fitted to empirical dynamic experimental data.

Recently, pressure measurements in small-scale batch cultivations were applied as a measure of the catabolic activity (Borch et al. 2026). This enables the cultivation of the entire microbial population in parallel throughout the entire cultivation cycle while quantifying catabolic activity. This work extends on the methodology applied by (Benthin et al. 1994), but apply pressure for catabolic rate measurements and online backscatter measurements to quantify anabolism and the biomass growth rate simultaneously. As illustrated in Table 1. Various metabolisms have very different gas-contribution ratios, which result in different slopes in the catabolic and anabolic visualisations, enabling identification of the present microbial metabolism at each timepoint and also when a single strain or the microbial community shifts between them.

**Table 1.** catabolic reaction stoichiometry. The overall stoichiometry of gas changes for various catabolic and anabolic reactions. Table from (Borch et al. 2026) reproduced with permission from the authors.

| Metabolism | Solubles | | | | | | | Gasses | | | | | Other | | | ΔGas |
| --- | --- | --- | --- | --- | --- | --- | --- | --- | --- | --- | --- | --- | --- | --- | --- | --- |
|  | Glucose | Lactate | Ethanol | Acetate | Methanol | Butyrate | Ammonia | Oxygen | CO<sub>2</sub> | CO | CH<sub>4</sub> | H<sub>2</sub> | Protein | Biomass | Water |  |
| Aerobic Glycolysis | -1 |  |  |  |  |  |  | -6 | 6 |  |  |  |  |  | 6 | 0 |
| Alcoholic Fermentation | -1 |  | 2 |  |  |  |  |  | 2 |  |  |  |  |  | 0 | 2 |
| Lactic Fermentation | -1 | 2 |  |  |  |  |  |  |  |  |  |  |  |  |  | 0 |
| Homoacetogenesis |  |  |  | 1 |  |  |  |  | -2 |  |  | -4 |  |  | 2 | -6 |
| Carboxydutrophic acetogenesis |  |  |  | 1 |  |  |  |  | 2 | -4 |  |  |  |  | -2 | -2 |
| Methyloctrophic acetogenesis |  |  |  | 3 | -4 |  |  |  | -2 |  |  |  |  |  | 2 | -2 |
| Hydrogenotrophic methanogenesis |  |  |  |  |  |  |  |  | -1 |  | 1 | -4 |  |  | 2 | -4 |
| Aceticlastic methanogenesis |  |  |  | -1 |  |  |  |  | 1 |  | 1 |  |  |  |  | 2 |
| Chain elongation |  |  | -6 | -4 |  | 5 |  |  |  |  |  | 2 |  |  | 4 | 2 |
| Ethanol oxidation |  |  | -1 | 1 |  |  |  |  |  |  |  | 2 |  |  | -1 | 2 |
| Autotrophic anabolism |  |  |  |  |  |  | -0.4 |  | -2 |  |  | -4.2 |  | 2 | 3 | -6.2 |
| Glycolytic anabolism | -1.1 |  |  |  |  |  | -1.2 |  | 0.4 |  |  |  |  | 6 | 2.7 | 0.4 |
| Proteolytic anabolism |  |  |  |  |  |  | 0.3 |  | 0.5 |  |  |  | -1 | 0.5 | -0.9 | 0.5 |

The aim is to map both catabolic and anabolic rates from simple, single, closed-batch cultivation onto a phase plane to visualise metabolism through an entire anaerobic cultivation cycle. Besides characterising the individual cultivations, a motivation and hypothesis were the potential generation of entire growth-cycle, non-disturbed, non-growth-enforced dynamic data sets. These could have implications for the fitting of dynamic metabolic models. Despite the strong practical suitability for combining online pressure measurement and backscatter for closed anaerobic sterile cultivation workflows, this approach has rarely been reported. We applied recently developed online pressure sensors (Borch, Kehr, et al. 2026). For this work, we further developed a plug-in for the Pioreactor system to enable simultaneous measurement of online pressure and backscatter in small-scale batch cultivations. First, we demonstrate this combined pressure-backscatter analysis during an anaerobic, high-sugar yeast cultivation. Secondly, we applied the framework to characterize a thermophilic syngas fermentation and showed how separate biomass and pressure trajectories reveal shifts between growth supported by yeast extract and two subsequent gas-based metabolic phases.

## 2 Materials and Methods

### 2.1 Pressure sensor plug-in for Pioreactor

The pressure sensors used were the Laerke.eu pressure sensor, recently developed in our lab (Borch, Kehr, et al. 2026). The sensors are designed to integrate with anaerobic, sterile workflows, using a needle to measure pressure through the rubber stoppers of vials or chemostats. For this work, we developed a software plug-in for the Pioreactor platform, a microbioreactor platform. The plug-in can be installed in the Pioreactor graphical interface through the Raspberry Pi repository: https://pypi.org/project/laerke-eu-pressure-reading-plugin/.

### 2.2 Yeast cultivation

We used Pioreactors v1.1 with a 60 ml N24 flask and a custom-designed butyl rubber stopper. Stirring at 800 rpm with a 12 x 3 mm magnetic stir bar. We used the laerke.eu Pioreactor plug-in version 1.0 and an I2C-connected Laerke.eu pressure sensors for each Pioreactor. The 120 ml vials were standard anaerobic vials with butyl rubber stoppers and crimp caps. Stirring at 350 rpm with 20 x 4 ml magnetic stir bars. The medium was prepared with dH_2_O, 20 g/L yeast extract, 80 g/L glucose, and adjusted to pH 6.5. The yeast was bought in a supermarket as a standard 50g wet baker’ s yeast and used the same day. The cultures were inoculated from an exponentially growing preculture (OD 6) to a starting OD of 0.05. The cultures were incubated at 34 °C.

### 2.3 Syngas fermentation

Anaerobic syngas batch vial cultivations were performed in 60 ml N24 vials with custom-made butyl rubber stoppers. The cultivation was controlled and monitored in the Pioreactor system v1.1 with: online backscatter; temperature control (60 °C), increasing the temperature a few degrees up from the incubator temperature set at 54 °C; magnetic stirring (12×3mm) at 800 rpm. Initial and final off-line OD600 samples were taken for approximate OD normalisation of the backscatter measurements. See pioreactor website, forum, and documentation for further description. The syngas composition used was (13% CO_2_, 30% CO, and 57% H_2_).

### 2.4 Acetogen media

The base components of the minimal media were dissolved in demineralized water (MiliQ) to reach the following concentrations: 1 g L^−1^ NaCl_4_, 0.4 g L^−1^ NH_4_Cl, 0.3 g L^−1^ KH_2_PO_4_. 2(n-morpholino) ethane sulfonic acid (MES) potassium salt was utilized as a buffer and added to a concentration of 18.7 g L^−1^, resazurin 4 mM. 10 mL L^−1^ of trace metal solution (modified from ATCC 1754 medium) with the following concentrations in mg L^−1^: Nitrilotriacetic acid 2000, MnSO_4_-H_2_O 1000, (NH_4_)_2_SO_4_-Fe(SO_4_)-6H_2_O 800, CoCl_2_-6H_2_O 200, ZnSO_4_-7H_2_0 200, CuCl_2_-2H2O 20, NiCl_2_-6H20 20, Na_2_MoO_4_-2H_2_O 20, Na_2_SeO_3_-5H_2_O 18, Na_2_WO_4_-2H_2_O 22, H_3_BO_3_ 10, AlCl_3_ 10.

Prior to use, 10 mL/L of stock solutions of vitamins, yeast extract, salts, and L-cysteine HCl were added to the base media to reach a final concetration of: L-cysteine 0.16 g L^−1^, Salts; 3 g L^−1^ MgCl_2_-6H_2_O, 0.05 g L^−1^ CaCl_2_-2H_2_O, 0.5 g L^−1^ yeast extract. Vitamin solution contained 0.012 g L^−1^ biotin (vitamin B7), 0.002 g L^−1^ folic acid, 0.010 g L^−1^ vitamin B6, 0.005 g L^−1^ vitamin B2, 0.010 g L^−1^ vitamin B1, 0.0004 g L^−1^ vitamin B12, 0.010 g L^−1^ vitamin B3, 0.005 g L^−1^ paminobenzoic acid, 0.030 g L^−1^ lipoic acid, 0.010 g L^−1^ Dpantothenic acid hemicalcium salt. The medium was prepared anaerobically and autoclaved at 140 °C for 25 minutes. For inoculation 5% v/v sample was used if nothing else is noted. The final pH of the medium is 6.8.

### 2.5 Syngas fermenting microbial community

The microbial community used as the inoculum for syngas fermentation was a novel enrichment developed for this work. It was based on a mixture of environmental samples: a biogas digestate, compost, and smoke scrubber liquid, collected from two industrial facilities, a coal-fired power plant, and a concrete factory, and enriched as previously described (Borch, Grimalt-Alemany, et al. 2026). For enrichment, online pressure sensors were used to minimize vial disturbances and facilitate subculturing timing. The environmental samples were stored at −20 °C until use. 20 ml of Milli-Q water was added along with the environmental samples; 20 ml of Ørsted coal power plant smoke scrubber slurry (50 °C), 20 ml of Solrød biogas slurry (45 °C), 15 ml of Aalborg Portland smoke scrubber (70 °C), and 15 ml of Solum biogas (35 °C). The mixture was sonicated for 20 seconds to homogenize it. It was then heat-shocked at 90 °C for 20 minutes in an autoclave. Afterward, the mixture was hand-shaken for 5 minutes to homogenise it, and the largest particles were allowed to sediment for 5 minutes, before being used as the inoculum source for the enrichment. After 4 rounds of cultivation, this enriched microbial community was used as an inoculum for the syngas cultivations.

### 2.6 Spectroscopic monitoring of cellular growth

The optical density (OD) was measured at 600 nm using a spectrophotometer, with MilliQ water as the blank (OD600). Samples with an OD ≥ 0.4 were diluted to stay within the linear detection range.

### 2.7 Chemical analysis

High-performance liquid chromatography (HPLC) analysis was used to analyse hydrolysate components and products. The supernatant was transferred to 2 ml autosampler vials. For the detection of carbohydrates and organic acids: column Aminex™ HPX87X ion exclusion (300 × 7.8 mm), flow 0.6 ml min^−1^, injection volume 20 μL, column temperature 65 °C, mobile phase 5 mM sulfuric acid in MilliQ, run duration 45 min.

### 2.8 Statistics and replication

Where shown in the figures, error bars represent variability across biological replicates and are reported as standard error of means (SEM) or as indicated in the corresponding figure captions.

## 3. Results

### 3.1 Metabolic phases in a baker’ s yeast cultivation

Baker’ s yeast was cultivated in closed anaerobic vials in rich medium containing 80 g/L glucose and 50 g/L yeast extract. Online pressure and backscatter data were analysed and plotted in R. The backscatter was blanked before inoculation, and the final backscatter was linearly scaled to the final offline OD measurement. This scaling is an approximation that introduces error because backscatter readings are not linearly correlated with biomass concentration; for precise analysis, several OD-backscatter measurements should be used. The cultivation was performed in both 120 ml and 60 ml vials, with 40 and 20 ml of medium, respectively, to enable comparison across experimental scales. The pressure curves for the two vial sizes aligned. For the Pioreactor setup, the pressure increased from 1 atm to ~8.5 bar absolute, ~7.5 bars overpressure. The 120 ml vials increased to ~8 bars overpressure. Linear plots of backscatter and pressure in Supplementary Figure 1. We applied a smoothing algorithm on the raw sensor data to extract the gas and backscatter rates. In Figure 1D,E, the measurements are plotted with the smoothed lines. The 120 ml vials had a slightly faster gas rate (Figure 1E), compared to the 60 ml. Pioreactors, but overall, they showed the same pattern.

**Figure 1.**
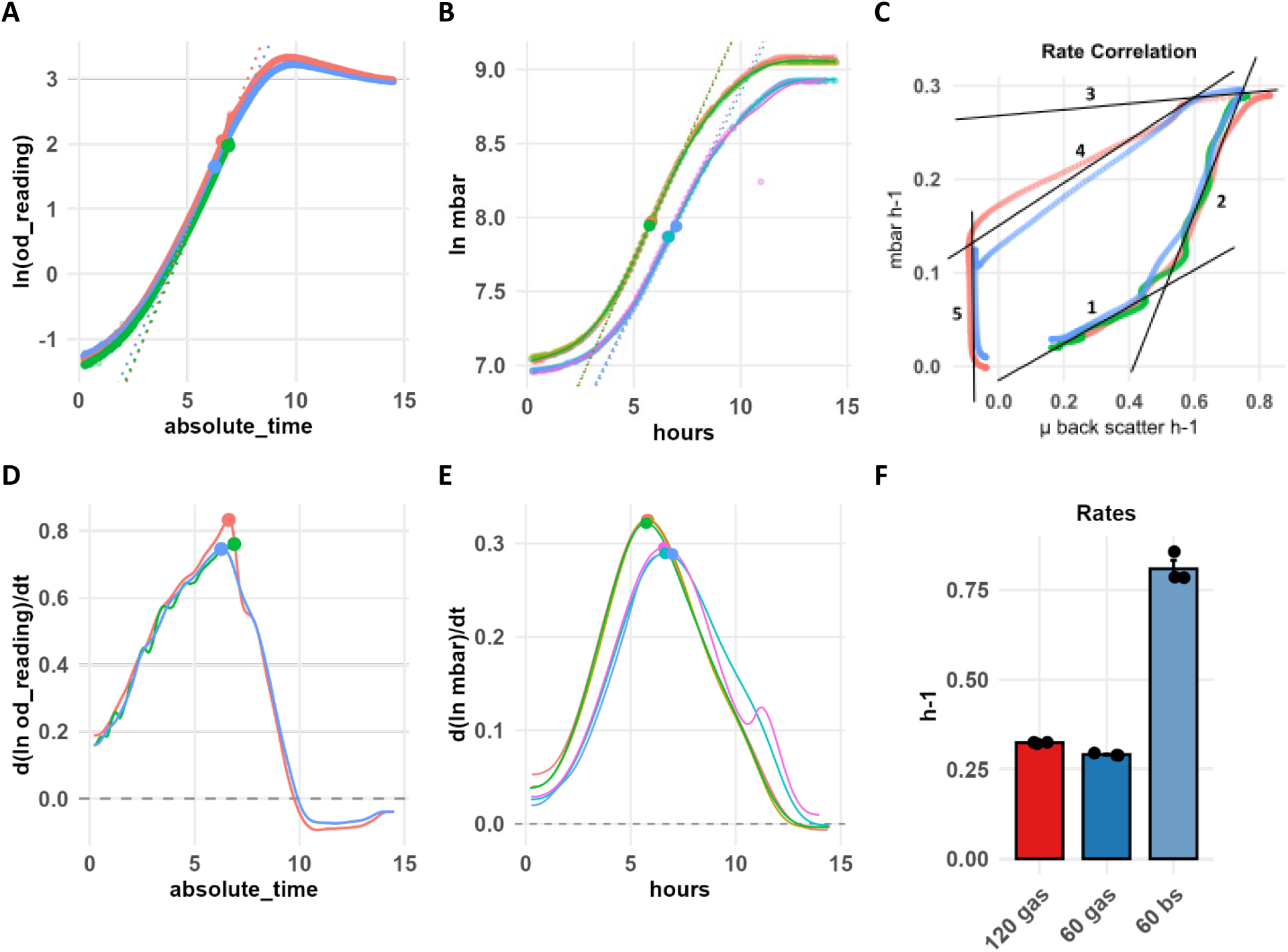
Backers’ yeast cultivation with online pressure and backscatter. Backscatter and pressure data for 60 ml condition and only pressure for 120 ml vials. A) Natural logarithm plot of the backscatter. B) Natural logarithm of mbar pressure data, the upper three curves are the 120 ml vials. C) phase plane plot of growth rate *μ* calculated from the backscatter vs pressure rate. 5 phases of cultivation are indicated by black lines and numbered. See main text for discussion. Rates estimated from a smooth spline of the ln plots. D) Growth rate *μ* h^−1^ (backscatter-based). E) Pressure rates. The three upper-left curves are for the 120 ml vials. F) Maximum pressure and growth rates from vials. Error bars are SEM of biological triplicates.

The backscatter rate and the pressure rate were plotted together. In this plot, 5 phases can be identified during the cultivation (Figure 1C) and we estimated the slope and intersection (Table 2). As a simplification, we set the gas contribution to anabolism to zero, enabling conversion of CO_2_ rates and yield to ATP-based rates and yield. The ATP requirement for biomass and maintenance in mmol/g DW was determined according to (Benthin et al. 1994). A key technical note is that the maximum growth (*μ*_max_) inferred from backscatter approached ~0.8 h^−1^ in this dataset. This is higher than the typical reported *μ*_max_values for *S. cerevisiae* under anaerobic conditions. This arises from the non-linear backscatter-to-biomass scaling applied at a single point and from the initial technical blanking of the backscatter. Furthermore, baker’ s yeast is often contaminated with lactobacillus strains, which have a *μ*_max_ of ~0.8 h^−1^ in nutrient-rich conditions. Methodologically and qualitatively, the analysis still holds, as visualization and evidence of metabolic correlations and shifts, that fit well with the known condition.

**Table 2.**
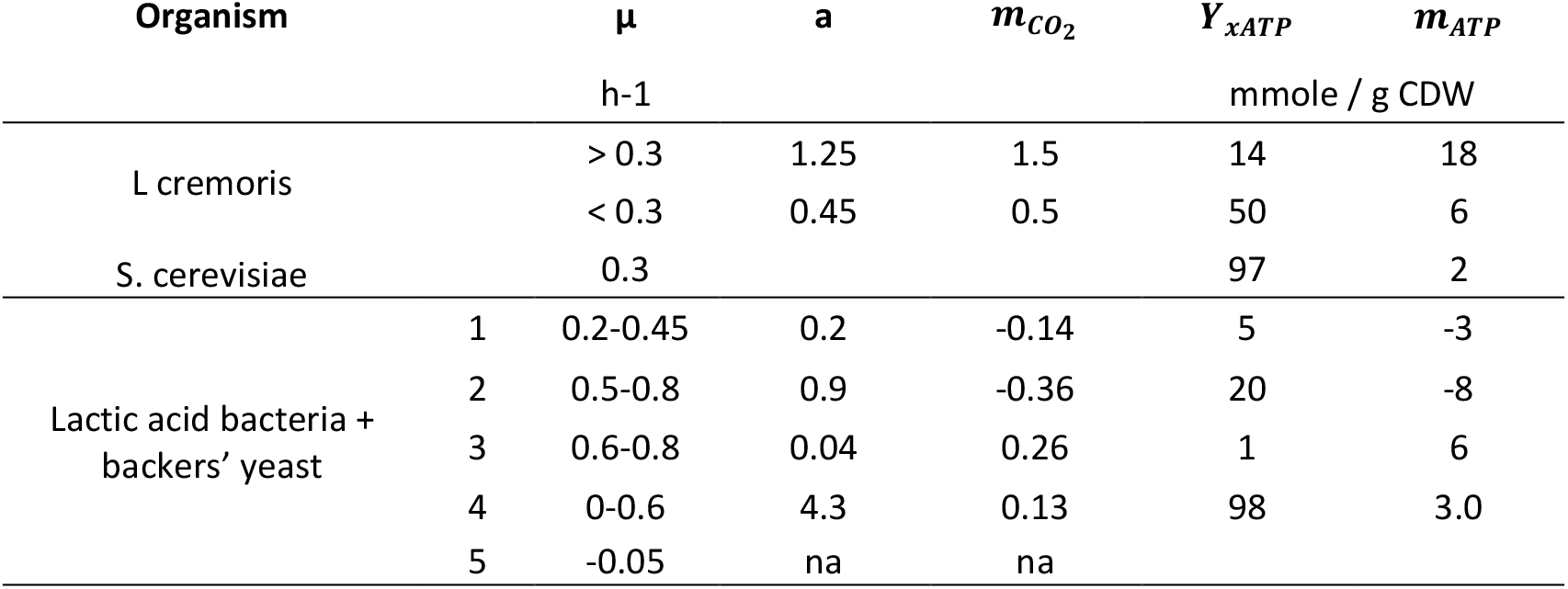
Growth and ATP correlation. Linear correlations between growth rates and ATP formation rates were either obtained through product formation rates or CO_2_ formation rates. Values from this work and (Benthin et al. 1994; Villadsen et al. 2011).

For the 5 cultivation phases, the following is noted: 1) During the initial phase of cultivation, the growth rate slowly increases, but pressure development is limited or negative. The slope is low, ~ 0.2. 2) During the second phase, the slope is ~ 0.9, indicating an even stoichiometric correlation between the gas formation and the growth rate. The two initial phases have *Y*_*xATP*_ below the theoretical minimum, and a negative maintenance value, illustrating catabolic underestimation measured through the CO_2_ formation. Several factors could contribute to this. In phase 1, there will be initial consumption of residual oxygen, and, based on the yeast aerobic anabolism stoichiometry (Table 1), this will generate approximately an equimolar amount of CO_2_. CO_2_ absorption in the media can also delay the pressure response and underestimate the final measured CO_2_. This is the case in this cultivation, due to the high final pressures reached, as also confirmed by heavy bubbling in the flasks, when they were degassed after the experiment. The CO_2_ formation rate can also be lowered due to a lower energy requirement. The medium contains a high concentration of nutrient-rich yeast extract. This lowers the energy requirement for biomass formation. Further energy/ATP can be generated by peptide hydrolysis in the cells, resulting in amino acid and proton symport efflux that leads to further energy generation (Benthin et al. 1994). 3) The third phase is nearly flat with a decreasing growth rate and a constant pressure formation rate. This slope and intersection is outside the range of previously reported experimental and theoretical values. We consider this a transition phase in which growth slows, while, for some time, the CO_2_ rate remains constant due to maintenance and CO_2_ equilibrating between the liquid and gas phases. 4) During the fourth phase, there is a decrease in gas and growth rates until growth completely stops and then drops below zero, indicating degradation or lysis of the cells. At *μ* = 0, the gas formation rate is 0.15 h^−1^, thus the non-growth-associated catabolic-maintenance rate. The ATP yield and maintenance of this phase is identical to previously obtained empirical values for the growth and non-growth associated maintenance of *S. cerevisiae* (Table 2). 5) During the fifth phase there is a constant biomass decay rate of ~ −0.07 h^−1^ while the gas rate continues, but gradually slows down, leading to continued increase of the total pressure. The final CO_2_ formation could be the maintenance of non-growing cells, and a small amounts of CO_2_ from the liquid phase balancing with the headspace pressure. Eventually, both the decay and pressure rates reach zero.

### 3.2 Metabolic phases in thermophilic microbial community syngas fermentation

In mixed community syngas (CO, H_2,_ CO_2_) fermentation, both carboxydotrophic (CO) and hydrogenotrophic (H_2_/CO_2_) metabolisms occur. CO has several inhibitory effects, in particular on hydrogenases (Imanishi et al. 2022), affecting the balance and transition between these metabolic modes. (Borch et al. 2026) demonstrated that even small amounts of yeast extract can increase the gas-uptake rates of CO-fermenting *C. autoethanogenum*, but here for the microbial communities, heterotrophic metabolism from hydrolysers in the community probably leads to additional formation of H_2_/CO_2_, as also demonstrated earlier (Borch, Grimalt-Alemany, et al. 2026). The CO metabolic stoichiometry (Table 1) and the inhibitory effect together could cause an initial constant total pressure. We earlier observed this in microbial community syngas batch fermentations (Borch, Kehr, et al. 2026). This experiment thus aimed to study biomass development during the initial stationary-pressure phase and to characterize the recently enriched syngas-fermenting microbial community. For this, the combination of online backscatter and pressure measurements was applied. We carried out a syngas fermentation in the Pioreactor setup (60 mL vials, 20 mL medium) using a standard anaerobic medium with 0.5 g/L yeast extract. The substrate was syngas (13% CO_2_, 30% CO, 57% H_2_) prepared at an absolute pressure of 2.5 bar (dry gas sparging at room temperature). The experiments were performed in technical triplicate, including a negative control without syngas to evaluate the effect of the yeast extract.

The sequential phases of the cultivation were distinguished and separated by identifying peaks in the gas consumption rate, growth rate, total gas transfer rate, and by visual inspection of the pressure and backscatter rate plot (Figure 2 A-H). The following phases were observed; numbers and letters refer to the phases and events as described and indicated (Table 3, Table 4, Figure 2). 0) no-growth: An initial phase was observed with no growth, and rapid and fluctuating pressure changes, until the vial temperatures stabilized. 1) Heterotrophic growth. At h 8 to 11, exponential growth (a) was observed both with and without syngas, though with syngas this rapid initial growth resulted in a higher backscatter than observed for the control, indicating a synergistic effect (Figure 2A). A slight increase in pressure was observed during heterotrophic growth. 2) Transition and exponential gas consumption. As yeast extract is depleted, the growth rate decreased rapidly, while the gas consumption rate increased as the community switched from heterotrophic to gas metabolism (h 11-22). A peak in gas consumption rate (h. 18) was followed by the highest growth rate at 22 hours of cultivation, likely due to growth on CO and H_2_/CO_2_ combined. 3) from h 22 to 41. Growth slows, and initially the pressure rate also slows, but then it rises to reach the second peak in gas consumption at h 38. Indicating a switch from CO to H_2_/CO_2_ metabolism. The increase in gas rate, with a drop in the growth rate, illustrates the stoichiometric difference in the number of gas molecules consumed between CO and H_2_/CO_2_ metabolism (Table 1). The change happens gradually and is not a sharp, abrupt shift. 4) hour 43 to 60. Gas-limited growth. The total gas transfer peaks at h 41, and then rapidly decreases. The cultures are from here gas-limited, and growth, although maintained, slows down as the last gases are consumed, and the gas consumption rate declines rapidly. 5) At h 60, a shift in the growth rate indicates a depletion of H_2_, and the halt of the hydrogenotrophic microbes in the community. Some growth is still observed, likely due to hydrolysis; thus, growth on the last substrates and decaying biomass, possibly slightly supported by headspace CO_2_, as some pressure consumption is retained. 6) At h 80, the gas-exchange completely stops, and both rates decrease to zero.

**Table 3.** Peaks in gas and growth rates. Timepoints and rates for observed peaks in gas consumption and growth during thermophilic syngas fermentation with a microbial community.

|  | Peaks / Events | hour |  |  |
| --- | --- | --- | --- | --- |
| a | $\mu_{\max}$, heterotrophic | 10 | 0.105 | h-1 |
| b | Gas consumption peak on CO | 18 | 0.36 | h-1 |
| c | $\mu_{\max}$, combined gases | 22 | 0.073 | h-1 |
| d | Gas consumption peak on H<sub>2</sub>/CO<sub>2</sub> | 38 | 0.32 | h-1 |
| e | Total maximum gas transfer | 41 | 67 | mbar h-1 |
| f | Growth rate shift, H<sub>2</sub> CO depletion | 60 | n.a. |  |

**Table 4.**
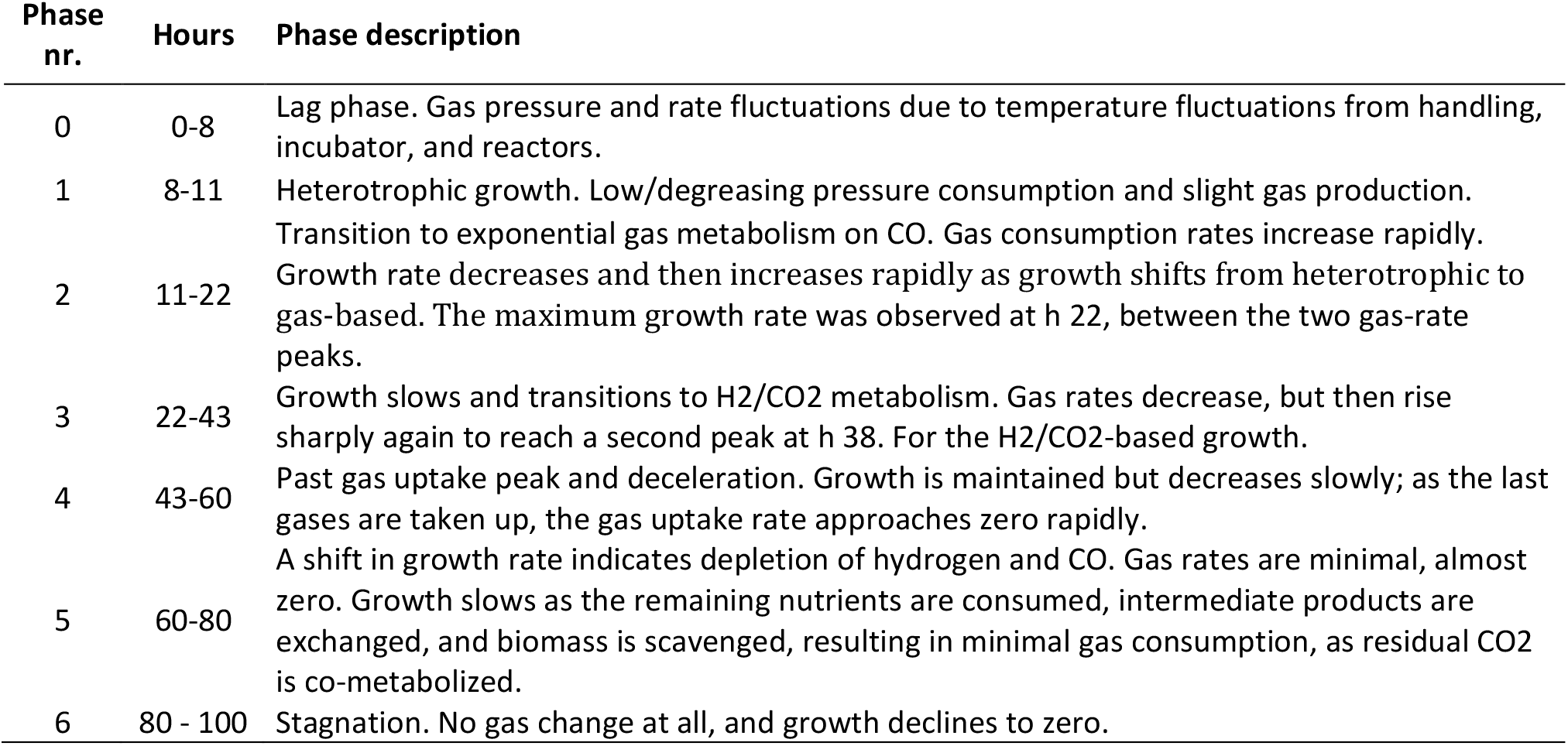
Metabolic phases in syngas fermentation. Description and duration of metabolic phases observed during the thermophilic syngas fermentation.

**Figure 2.**
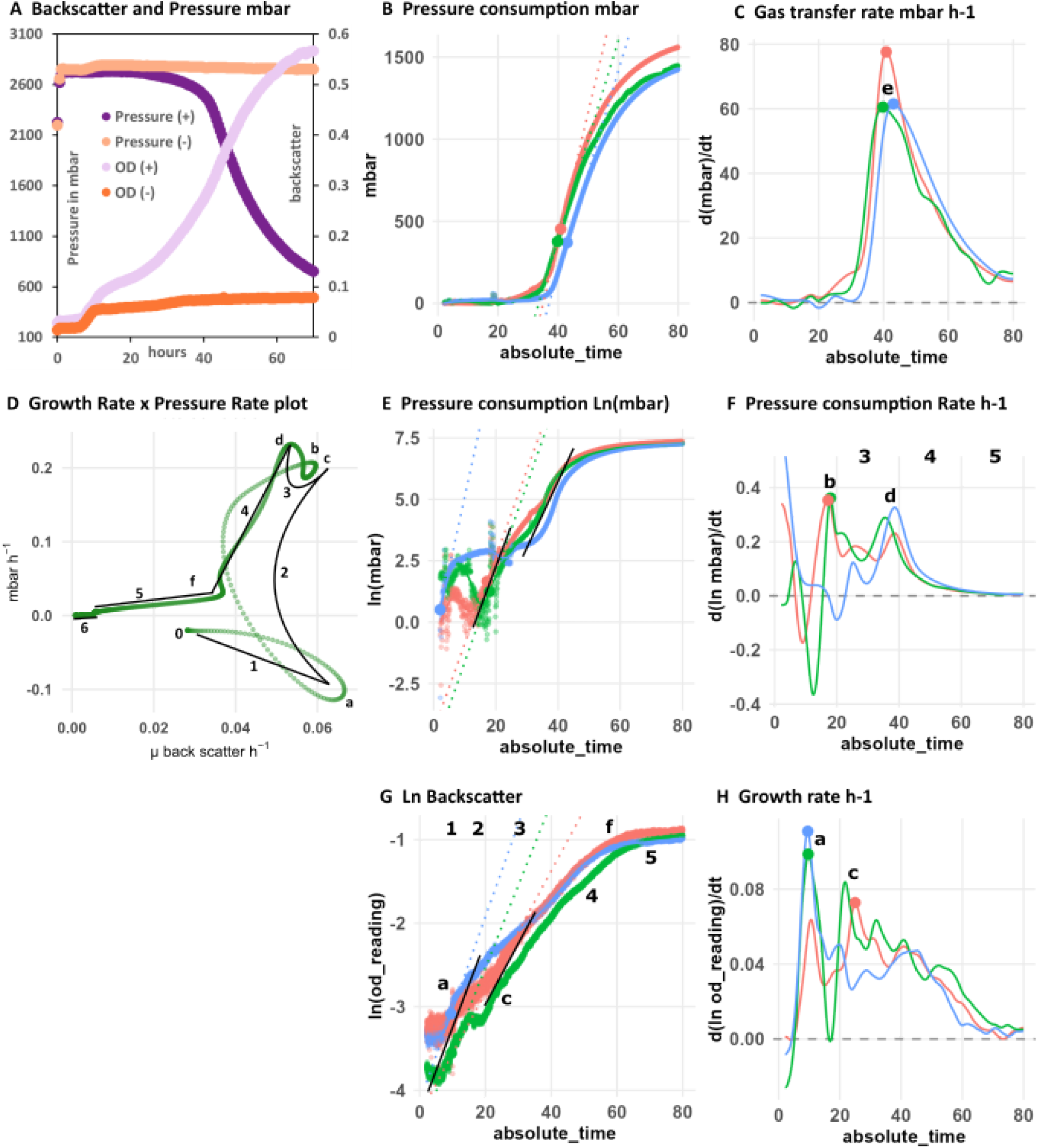
Pressure and backscatter data from microbial community thermophilic syngas fermentations. A) Average pressure (mbar) and optical density derived from backscatter measurements under syngas and negative control conditions. B) Total pressure consumption normalized to each cultivation’ s maximum measured pressure. C) Total gas transfer with a rapid increase after the initial CO-based gas-rate peak, and hence CO depletion (b). D) A rate-rate plot illustrating pressure and backscatter rates, with metabolic phases marked by straight lines and numbered, as detailed in the text and Table 5. E) Ln gas consumption plot, showing maximum values; the initial rate (h 2) is due to temperature fluctuations. The second and third phases correspond to exponential growth on CO and then increased H_2_/CO_2_ rates. F) Gas consumption rates starting with initial gas production from yeast extract metabolism, followed by a sharp rise to the CO-based rate peak (a) at h 18, then shifting to the H_2_/CO_2_ peak (d) at h 38. G) Ln backscatter plot indicating exponential growth phases: (a) early growth on yeast extract, (c) growth on CO, and a gradual decline in H_2_/CO_2_ growth (4), until hydrogen depletion causes growth to shift (f) to minimal levels.

The six phases described can be visually identified in the pressure and growth rate plot. All three replicates show the same overall pattern (Supplementary Figure 2), but they do not align, and for clarity, only a single vial cultivation is shown (Figure 2D). The curves do not align, as they are technical replicates, and do not experience the exact same temperature fluctuations during early cultivation, which greatly affect the estimated gas rates, as seen from the early phase 0-20h of Figure 2E. The pattern in the pressure-growth-rate plot is very dynamic, with no clear straight-line phases, as could be identified for the yeast cultivation. Instead, the microbial community cycles and gradually transitions between many metabolic phases. The bottom-right corner represents heterotrophic metabolism, characterized by high growth rates and little to no change in pressure. Then, the two gas metabolic phases are in the upper-right corner. Top right, the CO metabolism with most efficient growth in relation to gas consumption, and then the H_2_/CO_2_ metabolic phase, with a slightly lower growth rate and increased gas consumption rate. This aligns with the stoichiometry for these two metabolisms (Table 1).

The main activity occurred between approximately 8 and 75 hours. Yeast extract supported growth from around h 8, but, unlike the observation for pure *C. autoethanogenum* cultures (Borch et al. 2026) where an increased gas consumption was observed, here a slight increase in pressure was observed instead, probably due to CO_2_ formation from faster-growing hydrolysing strains (Borch et al. 2026). The maximum gas transfer is a short peak, not a sustained gas transfer-limited phase. To investigate mapped metabolic phases further, the gas composition could be changed, and co-feeding strategies to alleviate the CO inhibition and bottlenecks could be applied (Mann et al. 2020). All the identified phases are short, making it very difficult to collect enough intrusive samples to characterize each one. We also see indications that manual, intrusive sampling drastically alters growth trajectories and dynamics by disturbing the cultures. Thus, for microbial communities or situations in which growth shifts through several phases, this online pressure-growth-rate analysis method could provide a useful alternative or supplement to manual sampling. If samples of specific phases are desired, an initial experiment could map the individual phases, and samples could be collected for phases of interest in a subsequent experiment. This could be useful for quantifying gene expression (transcriptomics), protein levels (proteomics), or metabolites at specific time points (metabolomics).

## 4 Discussion

Analysis of simultaneous, online biomass measurements, combined with product and substrate rates, is widely used. (Benthin et al. 1994) demonstrated the strength of the simultaneous measurement of biomass and product formation rates. Several online biomass and product measurement tools are available beyond the off-gas analysis mentioned. Near-infrared (NIR) and Fourier-transformed mid-infrared (FT-MIR) have been applied for real-time monitoring of carbon balances in chemostats (Beuermann et al. 2012), and in a batch bioreactor, a circulating loop of both the liquid and gas phase provided non-sampling high-quality online data (FTIR, Ramen) (Metcalfe et al. 2020). Several online tools available for anaerobic digestion have also recently been reviewed (Cruz et al. 2021).

The key novelty of this study, along with the tools and methodology presented, is providing non-intrusive, simple, and inexpensive measurements of pressure for estimating the catabolic rate, and backscatter to determine biomass rates during a batch cultivation. An important limitation is the observation that the pressure measurements are very sensitive to temperature fluctuations, as previously discussed (Borch et al. 2026). This highlights the importance of non-disturbed, stable temperature conditions before inoculation to separate temperature effect from microbial effects.

## 5 Conclusion and perspectives

The work establishes a practical framework for catabolism-resolved microbial characterization in sealed batch vials by coupling high-resolution online pressure (gas exchange) with online backscatter/OD (biomass). We applied it to anaerobic batch cultivation and mapped the metabolic phases and shifts during the cultivation of yeast and a syngas-fermenting microbial community.

To expand on this work, further cultivations with different substrate and headspace gasses could be performed to strengthen the conclusions. Additionally, pure cultures of the strains identified in the microbial community could be grown to verify the findings presented here. A limitation of the setup is the temperature sensitivity, particularly during the early cultivation. The pressure-backscatter-rate approach introduced provides a framework for directly quantifying states and transition dynamics. Applications include: i) pinpointing the onset and timing of nutrient limitation, inhibition, and recovery. ii) mapping the effects of pulse feeding and intermittent gas supply. iii) screening pathway disruptions (such as inhibitors or knockouts) based on changes in rate-correlation. iv) real-time detection of co-feeding effects that shift catabolic allocation, alleviate bottlenecks, or modify yields. v) deriving constraints from slopes, intercepts, and thresholds to bound stoichiometric or kinetic models, which are not obtained from chemostats with forced growth or gas-sparging. vi) The entire dynamic rate-correlation could provide data and an interpretive framework for fitting or containing dynamic models.

## 6 Declarations

## Acknowledgments

The authors want to thank and appreciate Lars Woetmann Pedersen for his collaboration and development of the laerke.eu Pioreactor plug-in. They also thank all staff at FabLab RUC for their contribution, collaboration, support, and feedback on hardware and electronics development.

## Author Contributions and Information

Conceptualisation, hardware and method development, and first draft, MMB. Data acquisition and analysis, MMB, PK. All authors contributed to data interpretation, revision, comments, and approval of the final manuscript.

